# Conditional KCa3.1-Transgene Induction in Murine Skin Produces Pruritic Eczematous Dermatitis With Severe Epidermal Hyperplasia and Hyperkeratosis

**DOI:** 10.1101/759274

**Authors:** Javier Lozano-Gerona, Aida Oliván-Viguera, Pablo Delgado-Wicke, Vikrant Singh, Brandon M. Brown, Elena Tapia-Casellas, Esther Pueyo, Marta Sofía Valero, Ángel-Luis Garcia-Otín, Pilar Giraldo, Edgar Abarca-Lachen, Joaquín C. Surra, Jesús Osada, Kirk L. Hamilton, Siba P. Raychaudhuri, Miguel Marigil, Ángeles Juarranz, Heike Wulff, Hiroto Miura, Yolanda Gilaberte, Ralf Köhler

## Abstract

Ion channels have recently attracted attention as potential mediators of skin disease. Here, we explored the consequences of genetically encoded induction of the cell volume-regulating Ca^2+^-activated KCa3.1 channel (*Kcnn4*) for murine epidermal homeostasis. Doxycycline-treated mice harboring the KCa3.1+-transgene under the control of the reverse tetracycline-sensitive transactivator (rtTA) showed 800-fold channel overexpression above basal levels in the skin and solid KCa3.1-currents in keratinocytes. This overexpression resulted in epidermal spongiosis, progressive epidermal hyperplasia and hyperkeratosis, itch and ulcers. The condition was accompanied by production of the pro-proliferative and pro-inflammatory cytokines, IL-β1 (60-fold), IL-23 (34-fold), IL-6 (33-fold), and TNFα (26-fold) in the skin. Treatment of mice with the KCa3.1-selective blocker, Senicapoc, significantly suppressed spongiosis and hyperplasia, as well as induction of IL-β1 (−88%), IL-23 (−77%), and IL-6 (−90%). In conclusion, KCa3.1-induction in the epidermis caused expression of pro-proliferative cytokines leading to spongiosis, hyperplasia and hyperkeratosis. This skin condition resembles pathological features of eczematous dermatitis and identifies KCa3.1 as a regulator of epidermal homeostasis and spongiosis, and as a potential therapeutic target.

## INTRODUCTION

Ion channels have long been known to contribute to the pathophysiology of inflammatory, autoimmune [1], and proliferative diseases [2–4]. More recently, several calcium-permeable channels of the transient receptor potential family (TRP) and potassium channels have been found to be involved in skin conditions, such as melanoma [5], psoriasis [6], atopic dermatitis [7], Olmsted syndrome [8], and rosácea [9–11], suggesting the respective channels as potential treatment targets.

One of the K^+^ channels considered a skin ion channel is the intermediate-conductance Ca^2+^–activated K^+^ channel, KCa3.1, encoded by the *KCNN4*-gene [12–14]. Its calmodulin mediated activation produces K^+^ efflux and membrane hyperpolarization, thus serving its multiple biological functions such as erythrocyte volume decrease [15–16], hyperpolarization-driven Ca^2+^-influx, proliferation and cytokine production in T-cells [1, 17], migration and activation of macrophages/microglia [18–19], Cl^−^ and H_2_O secretion in epithelia [20–21], as well as endothelium-derived hyperpolarization-mediated vasodilation [22–23].

KCa3.1 induction has been implicated in several diseases states characterized by excessive cell proliferation and inflammation (For recent in-depth reviews of KCa3.1 in health and as drug target in disease see [1–2, 24]. For instance, induction of KCa3.1 was shown to regulate the phenotypic switch of fibroblasts and smooth muscle cells towards a dedifferentiated proliferative phenotype that promoted pathological organ remodelling in the lung, heart, and kidneys [25–30], as well as arterial neointima formation [18, 31–32]. In addition, high expression of KCa3.1 has been considered a marker of tumor progression for some cancers [3, 33–34].

KCa3.1, therefore, can be viewed as a possible driver of disease, and several small molecule inhibitors for this therapeutically attractive target have been developed and tested in pre-clinical animal models [35–36]. The inhibitor, ICA-17043 (Senicapoc), initially intended for the treatment of sickle cell anemia, has been found clinically safe [37] and is currently being considered for drug repurposing for stroke and Alzheimer’s disease, two conditions in which KCa3.1 contributes to the pathophysiology [19, 38].

However, concerning the skin, the physiological role of KCa3.1 and its pathophysiological significance for human skin disease is largely unexplored. So far, KCa3.1 protein expression and/or mRNA message have been found in rat epidermis [39], in human keratinocytes and melonama [5], where pharmacological inhibition of KCa3.1 has anti-proliferative efficacy in vitro. Yet, the role of KCa3.1 in the healthy or diseased epidermis remains elusive. Here, we hypothesized that epidermal KCa3.1, as a channel that controls cell volume, proliferation and chloride and water-secretion, is a regulator of epidermal homeostasis.

To test the hypothesis, we generated conditional KCa3.1-overexpressor mice (KCa3.1+) that harbor a murine *Kcnn4*-transgene under the control of a tetracycline response element (TRE) together with a rtTA-transgene [40] for DOX-inducible *Kcnn4*-transgene expression specifically in epithelia including the epidermis (Fig 1A) and found that KCa3.1-induction causes eczematous dermatitis characterized by intra-epidermal edema (spongiosis), epidermal hyperplasia and hyperkeratosis, severe itch, and ulcers.

**Figure 1:**
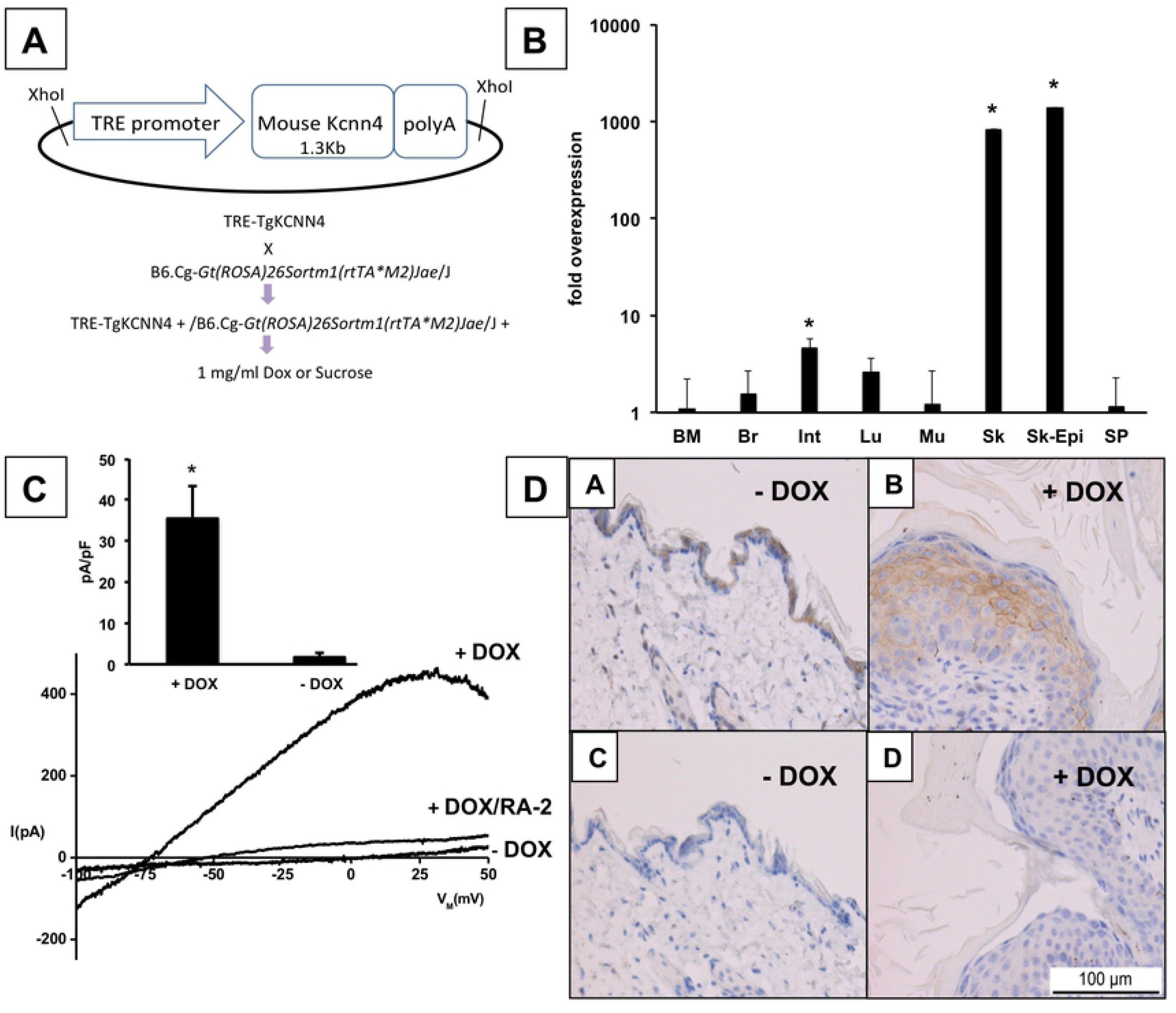
A) Plasmid construct for generation of *Kcnn4* transgenic mice and induction (gene product: KCa3.1) in epithelial tissues. B) Induction of KCa3.1 transgene expression by 2-weeks DOX-treatment over basal levels in various tissues as measured by qRT-PCR. Data (% of control (-DOX)) are given as means +/− SEM; *P<0.01, Student T test; BM, bone marrow (DOX, n=2; -DOX, n=2); Br, brain (DOX, n=4; -DOX, n=2); Int, small intestine (DOX, n=7; -DOX, n=6); skM, skeletal muscle (DOX, n=2; -DOX, n=2); Sk, skin (DOX, n=7; -DOX, n=6); Skin-Epi, skin epidermis (DOX, n=2; -DOX, n=2); Sp, spleen (DOX, n=4; -DOX, n=2); Lu, lung (DOX, n=11; -DOX, n=10). C) Whole-cell patch-clamp on freshly isolated keratinocytes from tail skin. Representative recordings of large KCa3.1 currents in keratinocytes (+DOX) from DOX-treated mice and currents in keratinocytes from untreated Ctrls (-DOX). Note: For an additional recording of small KCa3.1 currents in a keratinocyte from untreated Ctrl see Figure S1A. Inhibition of KCa3.1 currents by RA-2 at 1 μM. Inset: Summary data of KCa3.1-outward currents at a clamp potential of 0 mV. Data (pA/pF) are given as means +/− SEM (-DOX, n= 4; DOX n=5); *P<0.01, Student’s T test. D). Immune histochemical detection of KCa3.1 protein in the epidermis of DOX-treated and untreated mice (-DOX).

## MATERIAL & METHODS

### Transgenic Mic

Our TRE-Tg*Kcnn4* mice were generated at Unitech Co., Ltd. (Chiba, Japan). Briefly, a tetracycline-regulated *Kcnn4* expression construct was generated by subcloning PCR-amplified cDNA encoding the open reading frame of murine *Kcnn4* (gene ID16534)) into the pTRE-Tight expression vector (Clontech). The construct was verified by sequencing. The pTRE-Tight vector construct was cleaved with the restriction enzyme and injected into pronuclei of fertilized mouse oocytes of the C57BL/6J strain. The putative TRE-Tg*Kcnn4* founders obtained were genotyped by PCR with primers specific for the murine *Kcnn4* sequence. Two founders were crossed with wild type C57BL/6J mice to establish the F1 generation. One line was inbred over 2-3 generations and then crossed with B6.Cg-Gt(ROSA)26Sortm1(rtTA*M2)Jae/J + [40]. Routine genotyping was performed by using DNA from tail tips and PCR primers (see table 1) and the SuperHotTaq Master mix (BIORON GMBH, Germany); cycle program: 94 °C for 2 min, 35 cycles 94 °C for 20 sec, 56°C for 30 sec, 72 °C for 30 sec, and cooling to 10°C. PCR products were separated by gel electrophoresis (1.5% agarose). Pups being hemizygous for both trangenes were used for experimentation. All transgenic mice were generated and maintained within a specific pathogen free (SPF) barrier facility of Aragonese Center for Biomedical Research according to local and national regulations.

**Table 1:**
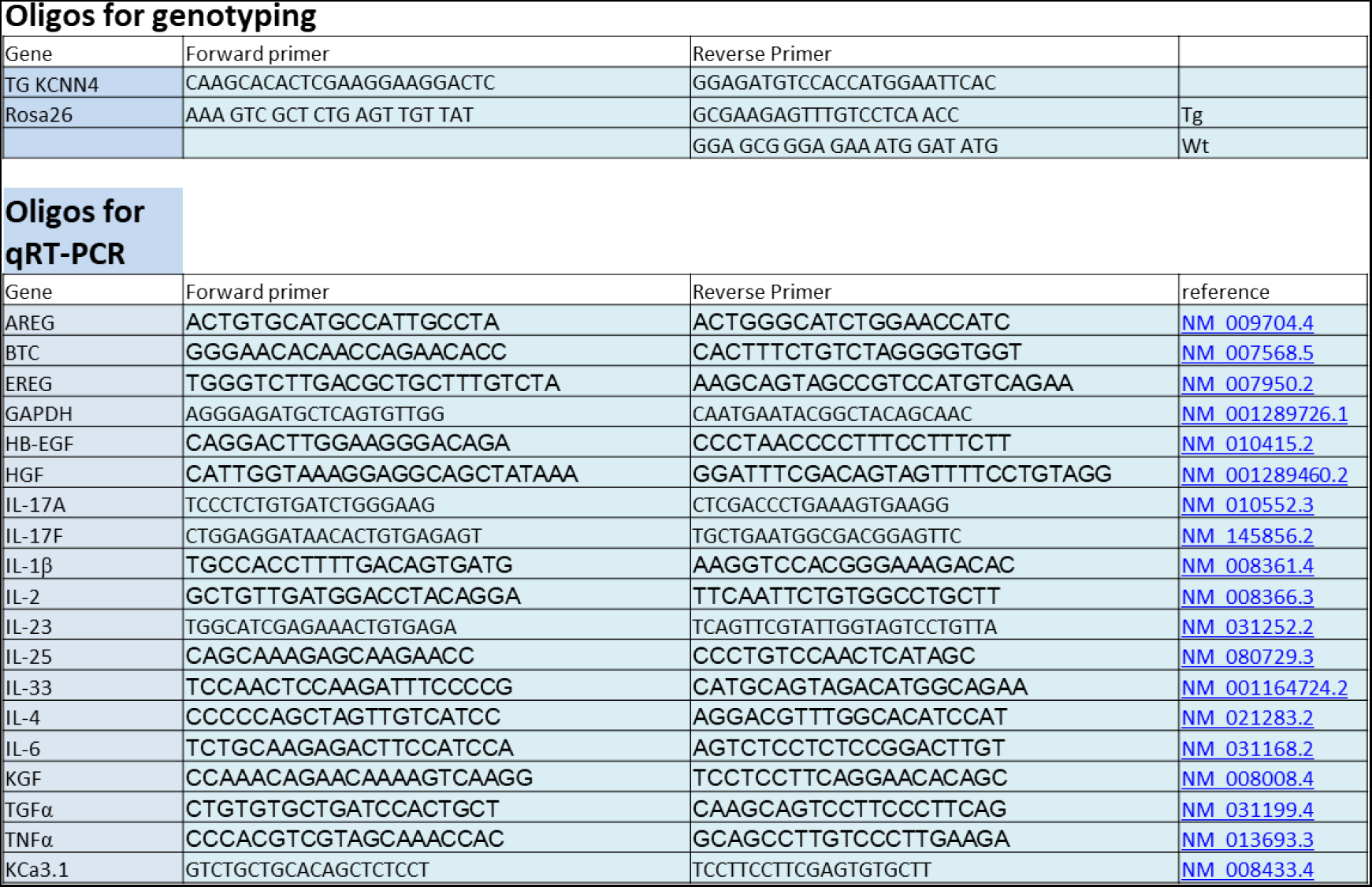
Primers for PCR and qRT-PCR

For transgene-induction over 1 or 2 weeks, doxycycline (DOX, Sigma) was added to the drinking water (1mg/ml) and water intake was monitored. Senicapoc was synthesized in-house as previously described. Senicapoc-medicated chow was prepared to administer a dose of 30 mg/kg/day. Photo- and video-documentation was routinely performed at the day of or the day prior to sacrifice. Organs and blood were collected after CO_2_ suffocation and stored on dry ice or fixed in neutral-buffered formaldehyde (4%) until further processing. All procedures were approved by the local Animal Ethics Committee (PI27/13; PI28/12; PI37/13/16; PI32/15) and in accordance with the ARRIVE guidelines.

### Isolation of Epidermal Keratinocyt

Tails were sterilized by short immersion in 90% ethanol and then stored in MEM-Earle with 20 mM HEPES until further processing. For separation of the skin from the underlying tissue, the skin was carefully cut open and removed from the base to the tip of the tail and cut into approximately 0.5 cm^2^ large pieces. Thereafter, pieces floated on a 0.25% trypsin/phosphate-buffered solution (PBS) overnight. For keratinocyte isolation, epidermis and dermis were carefully separated and the epidermis was cut into small pieces using a scalpel. Keratinocytes were dispersed by repeated passing through the tip of a cell culture pipette, seeded on coverslips in MEM Earle supplemented with 10% calf serum, and used for patch-clamp experiments within the next 3 hrs.

### Patch-Clamp Electrophysiology

Ca^2+^-activated K^+^ currents in murine keratinocytes were measured in the whole-cell configuration using an EPC10-USB amplifier (HEKA, Electronics, Lambrecht-Pfalz, Germany) and a pipette solution (intracellular) containing 1 μM Ca^2+^ free for channel activation (in mM): 140 KCl, 1 MgCl_2_, 2 EGTA, 1.71 CaCl_2_ (1 μM [Ca^2+^]_free_) and 5 HEPES (adjusted to pH 7.2 with KOH). The bath solution contained (in mM): 140 NaCl, 5 KCl, 1 MgSO_4_, 1 CaCl_2_, 10 glucose and 10 HEPES (adjusted to pH 7.4 with NaOH). For KCa3.1 inhibition, we applied 1,3-Phenylenebis(methylene)bis(3-fluoro-4-hydroxybenzoate (RA-2) at a concentration of 1 μM (n=2, experiments). Data acquisition and analysis was done with the Patch-Master program (HEKA). Ohmic leak conductance of up to 1 nS was subtracted where appropriate. We quantified outward currents at a potential of 0 mV. Membrane capacitance was 6 +/−1 pF (n=4) in keratinocytes from Dox-treated mice and 7 +/−2 pF (n=5) in keratinocytes from non-treated mice.

### Histology

Samples were fixed in 4% formaldehyde for at least 24 hrs and then transferred to 60% ethanol. Thereafter samples were imbedded in paraffin and cut into 4 μm-thick sections. Sections were stained with hematoxylin and eosin. Scoring: For quantifying and comparing the skin pathology, skin sections were scored for grade of hyperplasia, and hyperkeratosis by a pathologist (MM) and for grade of intra-epidermal edema by two investigators (RK/KLH), independently and in a blinded fashion. We used the following in-house developed scoring system: Mild: ≥0.5 <1; Moderate: ≥1<2; Severe: ≥2. Individual scores for intra-epidermal edema did not vary by more than 0.5 and were averaged.

### Immunohistochemistry and TUNEL Assay

For KCa3.1, tissue samples were fixed in 4% neutral buffered formaldehyde solution and embedded in paraffin. 2.5 μm-thick sections were cut with a rotation microtome (Leica RM2255). Slides were air dried at 37°C overnight, were de-paraffinized in xylene for 10 min, and then rehydrated. After rehydration, epitope retrieval was carried out using the PT Link (Dako) at 95°C for 20 min in a high pH buffer (Dako Antigen retrieval, high pH). Endogenous peroxidase was blocked using the EnVision FLEX Peroxidase-Blocking kit followed by two washes for 5 min each (Dako wash buffer). Sections were incubated for 60 min with a rabbit anti-KCNN4 primary antibody (AV35098, Sigma-Aldrich) at 1/2000 dilution followed by two washes. Signal amplification was done using the ImmPRESS™ Excel Amplified HRP Polymer Staining Kit (Vector Laboratories). After 3 wash steps (Dako wash buffer, 5 min each). Sections were incubated with 3,3’-diaminobenzidine (DAB) for 10 min and counterstained with hematoxylin.

Proliferating cell nuclear antigen (PCNA) and Caspase 3 (CASP3): 5 μm-thick tissue sections were placed on silanated slides. After de-paraffinization, endogenous peroxidase was quenched by immersing the samples in methanol containing 0.03% hydrogen peroxide. Heat antigen retrieval was performed in pH 6.5 10 μM citrate-based buffered solution (Dako). In order to prevent non-specific binding, samples were incubated for 2 h with Protein Block (Dako). Samples were incubated overnight at 4°C with mouse and rabbit monoclonal antibodies against PCNA (Cell Signaling, 1:100) and Cleaved-CASP3 (Calbiochem, 1:100), respectively, followed by incubation with an anti-mouse or anti-rabbit polymer-based Ig coupled with peroxidase (Cell Signaling) for 30 min at RT. Then, sections were incubated with 3,3’-diaminobenzidine (DAB) and counterstained with hematoxylin, and studied under an Olympus BX-61 microscope.

Apoptosis was determined by using the TUNEL (Terminal deoxynucleotide transferase mediated X-dUTP nick end labeling) assay. 5 μm-thick sections were de-paraffinized and rehydrated. Then, sections were treated with Proteinase K (20 μg ml– 1, 15 min, at RT) and rinsed twice with phosphate buffered saline (PBS). For detection of apoptotic nuclei, sections were incubated for 1 hrs at 37°C in the dark with the *in situ* Cell Death Detection Kit (Roche) according to the manufacturer’s instructions. After two washes with PBS, sections were mounted in ProLong^®^ with DAPI (Life technologies). Fluorescence-microscopy was performed with an Olympus BX-61 epi-fluorescence microscope equipped with filter sets for fluorescence microscopy: ultraviolet (UV, 365 nm, exciting filter UG-1) and blue (450-490 nm, exciting filter BP 490). Photographs were taken with a digital Olympus CCD DP70 camera.

### RNA Isolation, Reverse Transcription, and Quantitative RT-PCR

Depilated skin of the neck and other organs were placed into 1 ml TriReagent (Sigma, Saint Louis, Missouri, USA) and stored at −80°C. Samples were homogenized with a T10 basic ULTRA-TURRAX (IKA, Staufen, Germany) at 4°C.

Total RNA was isolated with the TriReagent following the manufacturer’s protocol, and further purified using RNA Clean-up and Concentration-Micro-Elute kit (Norgen Biotek, Thorold, Canada). Genomic DNA was digested using the Ambion DNA-free kit (Invitrogen, Carlsbad, California, USA). Quantity and purity of extracted RNA were determined by spectrophotometry (NanoDrop1000, Thermofisher, Waltham, MA) and stored at −80°C for later use. Integrity of RNA samples and successful digestion of genomic DNA were verified by gel electrophoresis. Reverse transcription was performed with 600 ng of total RNA by using the Super Script III reverse transcriptase (Invitrogen, Carlsbad, California, USA) and random hexamers following the manufacturer’s protocol.

cDNA obtained from 10 ng of total RNA was amplified in triplicates using the SYBR Select Master Mix and a StepOnePlus Real-Time PCR system (Applied Biosystems, Foster City, California, USA) using the following cycle protocol: 95°C, 15 s and 60°C, 60 s repeated for 40 cycles. As final step, a melting curve analysis was carried out to verify correct amplification. The primers are given in table 1. Data were analyzed with LinReg PCR software and gene expression levels relative to *Gapdh* expression as reference gene and normalized to control were calculated using the formula: % *of Gapdh* =*Efficiency^Cq(Gapdh)−Cq(GOI)^×100*. The values were used to calculate ratio values (DOX/-DOX; Senicapoc/Ctrl) given in graphs.

### LC-MS Analysis

A 10 mM stock solution of Senicapoc (from in-house organic synthesis) was prepared by dissolving 6.4 mg Senicapoc in 2 ml acetonitrile. Working standard solutions were obtained by diluting the stock solution with acetonitrile.

Preparation of plasma samples: Commercial SPE cartridges (Hypersep C18, 100 mg, 1 ml) were purchased from Thermo Scientific (Houston, TX, U.S.A). Before extraction, cartridges were conditioned with acetonitrile, 2 × 1 ml, followed by water, 2-times × 1 ml. After loading the SPE cartridges with plasma samples, they were washed successively with 1 ml each of 20% and 40% acetonitrile in water followed by elution with 2 ml of acetonitrile. Elute fractions were collected and evaporated to dryness, under a constant flow of air, using a PIERCE Reacti-Vap^™^ III evaporator (PIERCE, Il, USA). The residues were reconstituted using 200 μl acetonitrile and were used for LC-MS analysis.

Preparation of skin samples: A 100 mg of depilated skin sample was homogenized thoroughly in 2.0 ml of acetonitrile in a gentleMACS™ M tube using a gentleMACS^™^ Dissociator (Miltenyi Biotec Inc., CA, USA). Each sample was subjected to three cycles of the preprogrammed homogenization protocol Protein_01.01 (a 55 s homogenization cycle with varying speeds and directions of rotation). Homogenized samples were centrifuged for 10 min at 4000 rpm. Each supernatant was collected in a 4 ml glass vial and was evaporated to dryness, under a constant flow of air, as described above. The residues were reconstituted in 100 μl acetonitrile and were used for LC-MS analysis.

LC/MS analysis was performed with a Waters Acquity UPLC (Waters, NY, USA) equipped with a Acquity UPLC BEH 1.7 μM C-18 column (Waters, New York, NY) interfaced to a TSQ Quantum Access Max mass spectrometer (MS) (Thermo Fisher Scientific, Waltham, MA, USA). The isocratic mobile phase consisted of 80% acetonitrile and 20% water, both containing 0.1% formic acid with a flow rate of 0.25 ml per minute. Under these conditions, Senicapoc had a retention time of 0.80 minute.

Using Heated electrospray ionization source (HESI II) in positive ion mode, capillary temperature 250 °C, vaporizer temperature: 30°C, spray voltage 3500 V, sheath gas pressure (N_2_) 60 units, Senicapoc was analyzed by the selective reaction monitoring (SRM) transition of its molecular ion peak 324.09 (M+1) into 228.07, 200.07,183.11 and 122.18 *m/z*. An 8-point calibration curve from 50 nM to 10 μM concentration range was used for quantification.

### Statistics

Data in Text and graphs are means +/− standard error of the mean (SEM), if not stated otherwise. If not otherwise stated, we used the unpaired Student’s T Test (two-tailed) for comparison of data sets. The significance level was set to a P value of <0.05.

## RESULTS

### KCa3.1 Induction in Skin

Doxycycline(DOX)-treatment with 1 mg/ml in drinking water produced a 833-fold KCa3.1-overexpression in the skin, 1378-fold KCa3.1-overexpression, particularly, in the epidermis, and a 46-fold overexpressing in the intestine (Fig 1 B). In bone marrow, brain, lung, skeletal muscle, spleen, KCa3.1-mRNA levels were similar to untreated mice.

Patch-clamp electrophysiology studies in isolated keratinocytes revealed a strong induction of KCa3.1-function in DOX-treated KCa3.1+ mice. The KCa3.1-currents demonstrated the typical fingerprint characteristics of KCa3.1, which were activation by 1 μM Ca^2+^, voltage-independent, and mildly inwardly-rectifying. RA-2 (1 μM), a negative-gating modulator of KCa2/3 channels [41], blocked the KCa3.1 current. KCa3.1-induction was not found in untreated mice and KCa3.1 outward-currents were small or difficult to discriminate from background currents in most cells (Fig 1 C). In fact, we saw a clearly distinguishable KCa3.1 current in only 1 of 5 cells (Fig S1A).

We examined the KCa3.1-protein expression by immune histology (Fig 1 D) and identified weak immune reactivity in the thin epidermal layer of untreated mice and appreciable immune reactivity in the thicker epidermal layer of DOX-treated mice (see following paragraphs).

Together, these findings demonstrate strong induction of KCa3.1-transgene expression and function in the skin and, particularly, in the epidermis.

### Skin Phenotype and Behavioral and Systemic Alterations in DOX-Treated KCa3.1+ Mic

The DOX-treated KCa3.1+ mice of both sexes appeared normal during the 1^st^ week of the treatment and then developed progressive skin pathology with generalized piloerection, intense scratching behavior (S Movie 1), scaly skin patches, and ulcerative lesions, mainly visible in the neck areas, chest, and ears (Fig 2), all sites where mice groom frequently.

**Figure 2:**
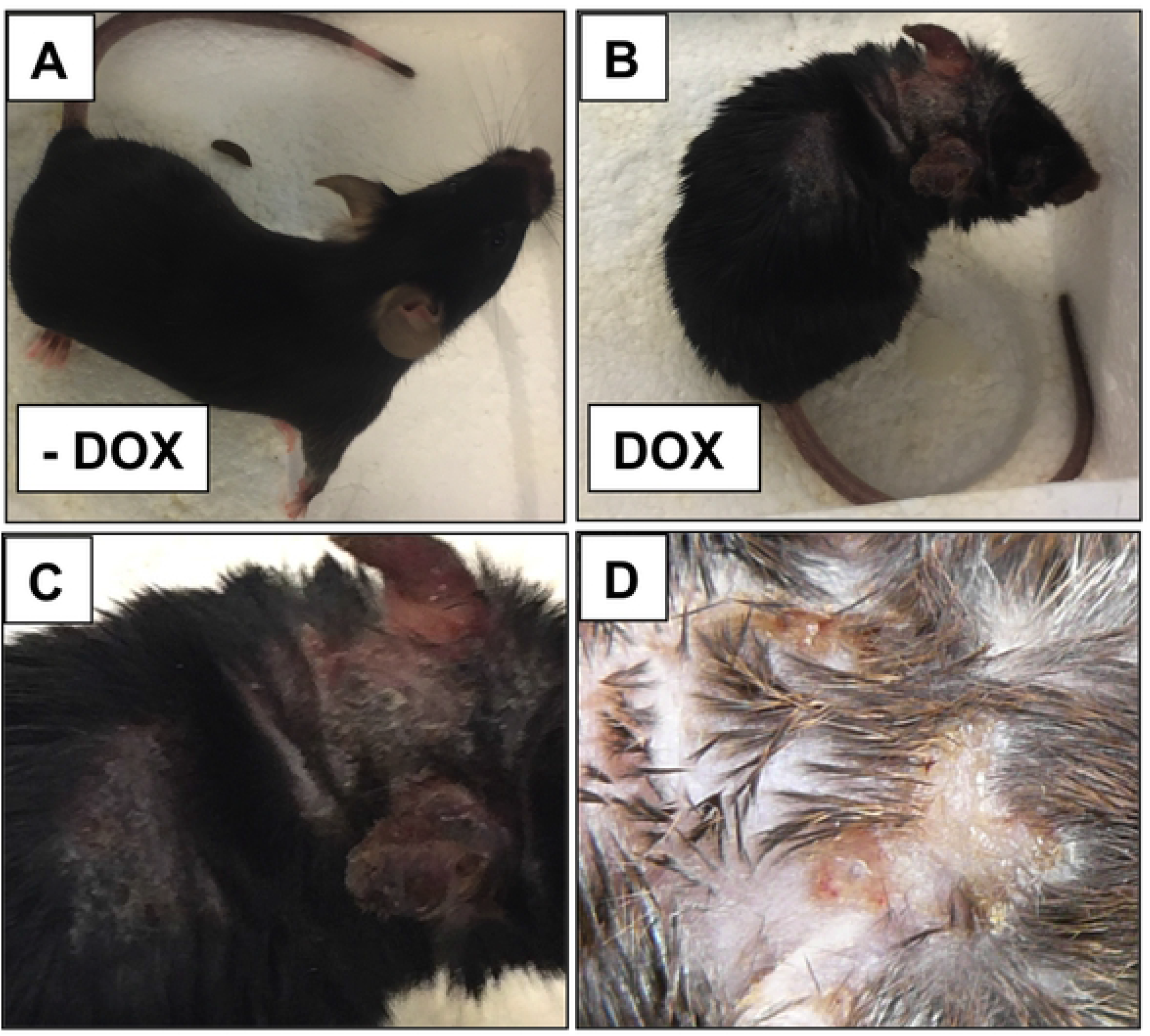
Macroscopic skin pathology DOX-treated KCa3.1+ mice. Photographs of A) untreated Ctrl (-DOX), B) DOX-treated KCa3.1+, C) higher magnification of the neck shown in B, D) patchy erythematous and scaly skin with ulcerative areas of a DOX-treated mouse. Note: Videos of DOX-treated mice showing severe scratching behavior and of Ctrls are found in the supplement.

Histology (Fig 3 A to D) of neck skin showed moderate to severe hyperplasia of the epidermis, affecting also the hair follicles, and substantial hyperkeratosis. In the epidermis of the DOX-treated mice, we found foci of moderate intra-epidermal edema (Fig 3 D), similar to spongiosis, which is characteristic of eczematous dermatitis [42]. The phenotype was completely reversible within 6 weeks when DOX was removed after 9 days of treatment (see S1B Fig).

**Figure 3:**
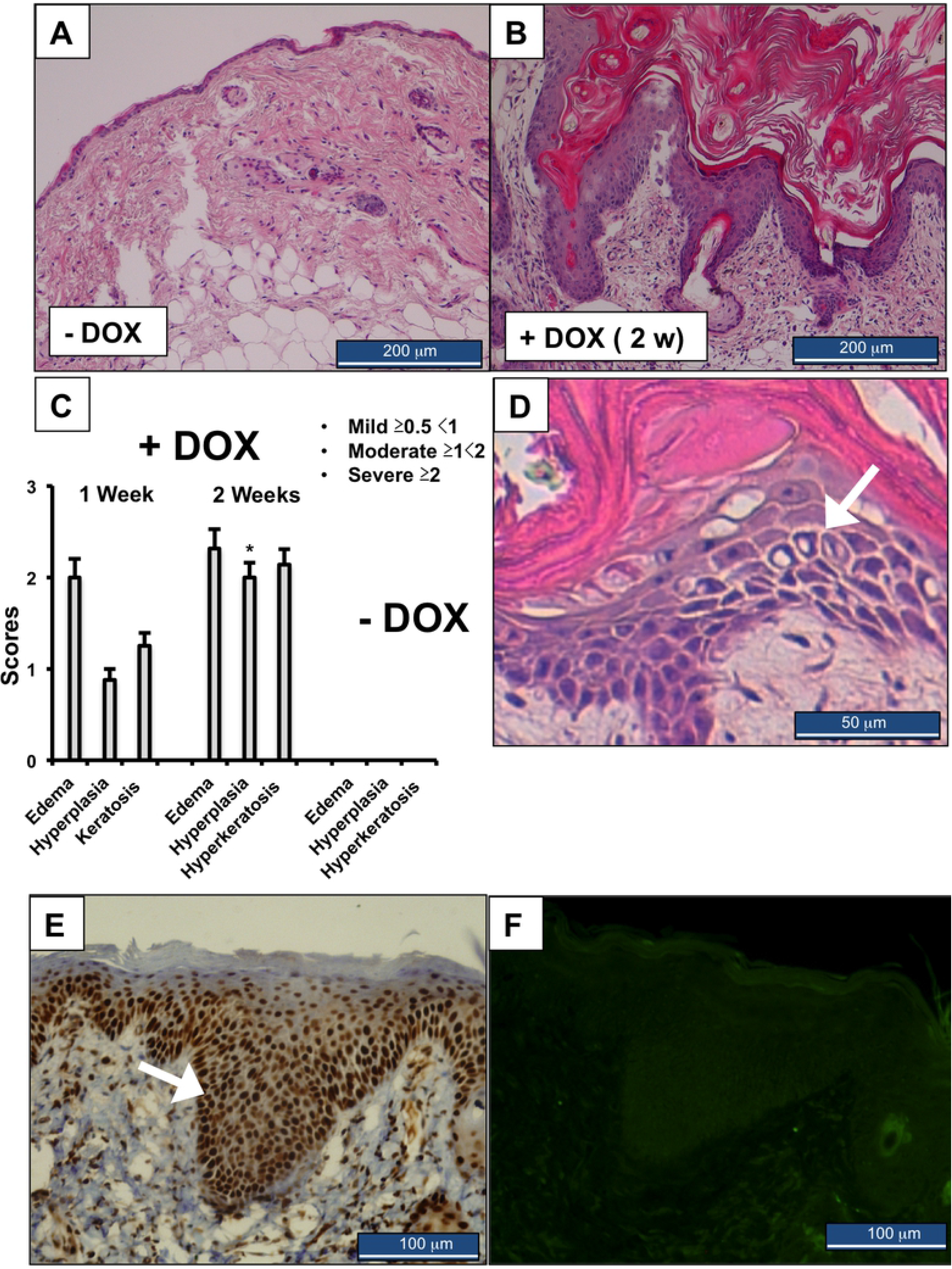
Histological evaluation of skin pathology in KCa3.1+ mice. H&E-stained sections of normal skin of an untreated mouse (A) and a skin of a DOX-treated KCa3.1+ (B) with severe hyperplasia and hyperkeratosis. C) Summary of pathology scores. Data are given as means +/− SEM, n=4 (1 week DOX), n=26 (2 weeks DOX), n=15 (Ctrls); *P < 0.05 vs. 1 week DOX, Student’s T test. Note that the scores for Ctrl skin are 0. D) Higher magnification of the hyperplastic epidermis of DOX-treated KCa3.1+. Note the presence of intra-epidermal edema with enlarged intra-cellular space (indicated by white arrow). E) Immune histological stains of the proliferation marker, PCNA, in the hyperplastic epidermis of DOX-treated KCa3.1+. Note the intense staining of the basal layer (white arrow) that becomes weaker when approaching the stratum corneum (representative image from 3 mice). F). The TUNEL assay detected no apoptotic keratinocytes in the hyperplastic epidermis.

At an earlier time point (1-week DOX), we recorded the same degree of intra-epidermal edema and an early stage of hyperplasia and hyperkeratosis (Fig 3 C and S1C Fig). Yet, at first sight, the mice appeared healthy. Concerning controls, a 2-week DOX-treatment of the inducer strain (R26-rtTA-M2) or of *Kcnn4*-transgene-harbouring mice lacking the rtTA induced no observable skin pathology (S1D Fig).

Immune histology on KCa3.1+ skins showed high-expression of the proliferating cell nuclear antigen (PCNA), a marker of cell proliferation, in the hyperplastic epidermal layer of 2-week DOX-treated mice (Fig 3 E and S1E Fig). In the untreated control animals, uniform immune reactivity was observed in the nuclei of keratinocytes lining the basal epidermal layer. We also stained for CASP3, another marker of apoptosis, and did not find induction of CASP3 in the hyperplastic epidermal layer or in the dermis, with the exception of ulcerative sites (S1E Fig). No CASP3 immune reactivity was seen in keratinocytes of the epidermis of untreated controls (S1F Fig). Apoptosis as measured by TUNEL was not found at hyperplastic sites (Fig 3 F), but at wound areas with substantial tissue destruction (S1E Fig).

Together, these data demonstrate that induction of KCa3.1 produces moderate intra-dermal edema (sub-acute spongiosis) and drives keratinocyte proliferation, but does not cause generalized cell toxicity and cell death. The localized epidermal damage and ulcers are likely the result of the intense scratching behavior.

### Alterations of Cytokine Expression Profile in Skin

To shed light on the mechanisms, by which KCa3.1-induction in the keratinocytes produced spongiosis and epidermal hyperplasia, we measured the mRNA-expression of several pro-proliferative cytokines that are known to promote keratinocyte proliferation in an auto-stimulatory or autacoid fashion [43] (Fig 4). It is worth mentioning that KCa3.1-KO and/or inhibition have been shown to reduce IL-β1 and TNFα levels in activated microglia after cerebral infarction [19].

**Figure 4:**
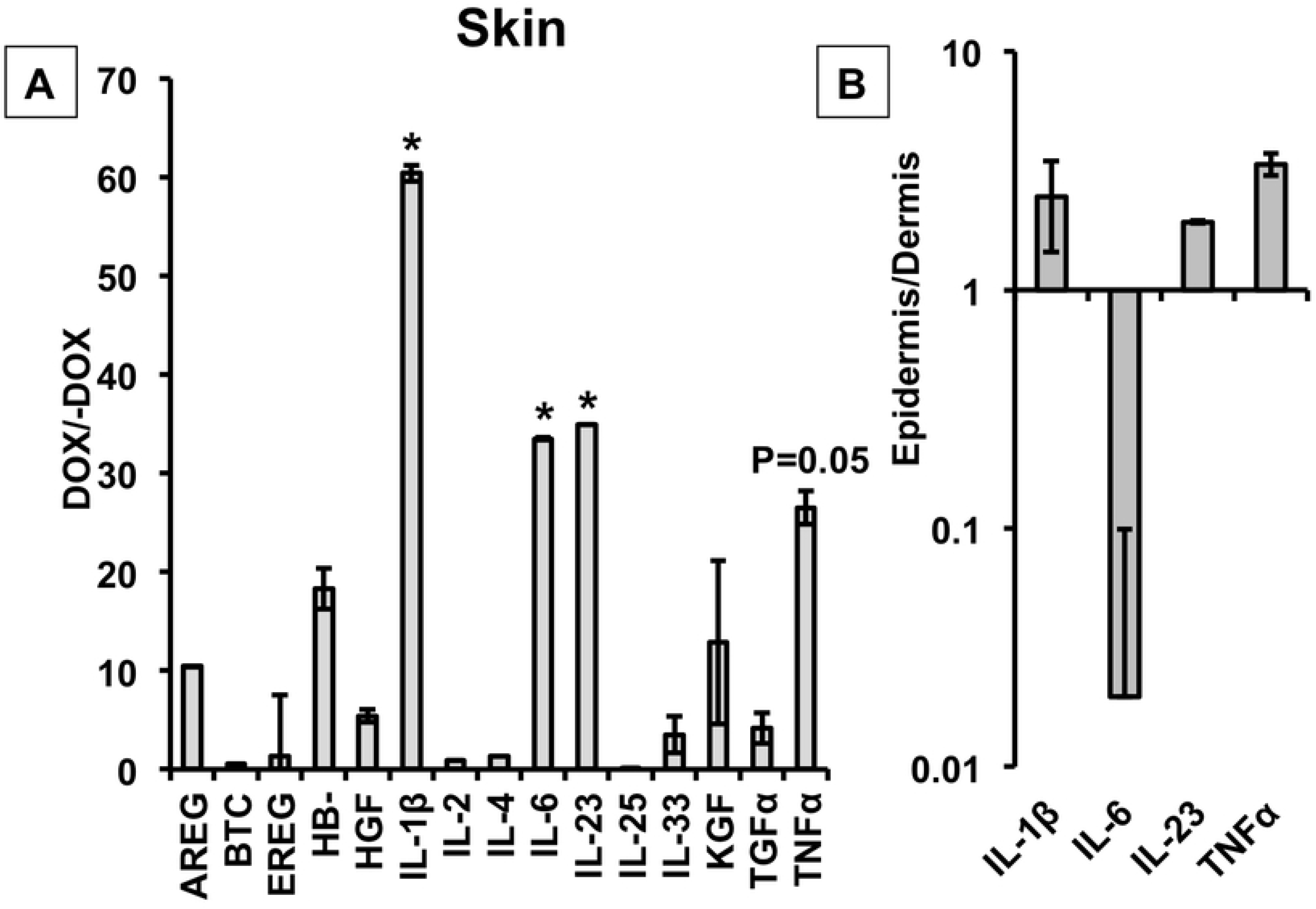
Cytokine mRNA-expression profile. (A) Alterations of cytokine mRNA-expression profile in DOX-treated KCa3.1+ mice (DOX/-DOX). (B) Cytokines with higher expression in the epidermis than dermis (epidermis/dermis). AREG, amphiregulin (DOX, n=2; -DOX, n=2); BTC, betacellulin (DOX, n=2; -DOX, n=2); EREG, epiregulin (dox, n=5; -DOX, n=5); HB-EGF, Heparin-binding EGF-like growth factor (DOXx, n=5; -DOX, n=5); HGF, Hepatocyte growth factor (DOX, n=5; -DOX, n=5), Interleukin(IL)-β1 (DOX, n=7; -DOX, n=7),IL-2 (DOX, n=2; -DOX, n=2), IL-4 (DOX, n=2; -DOX, n=2), IL-6 (DOX, n=7; -DOX, n=7), IL-23 (DOX, n=2; -DOX, n=6), IL-25 (DOX, n=2; -DOX, n=2), IL-33 (DOX, n=2; -DOX, n=2); KGF, keratinocyte growth factor (DOX, n=2; -DOX, n=2); TGFα, transforming growth factor α (DOX, n=5; -DOX, n=5); TNFα; tumor necrosis factor-α (DOX, n=5; -DOX, n=5). Data (DOX/-DOX; Epidermis/Dermis)) are given as means +/− SEM; *P<0.05, Student’s T test.

We found strongly increased mRNA expression of IL-β1 (60-fold), IL-23 (34-fold), IL-6 (33-fold), and TNFα (26-fold). Expression levels of HGF and TGFα as well as a series of other cytokines and growth factors, of which some are expressed by keratinocytes (amphiregulin, betacellulin, epiregulin, IL-2, IL-4, IL-25, IL-33, and keratinocyte growth factor, for review see [43], were not significantly altered (Fig 4 A). T-cell specific IL-17A and IL-17F mRNA expression was not detectable (data not shown).

The induction of IL-β1, IL-23, and TNFα expression was particularly high in the epidermal layer, although not statistically different from levels in the dermis. Yet, these data demonstrate strong cytokine induction in keratinocytes (Fig 4 B).

### Systemic KCa3.1 Channel Blockade Reduces Skin Pathology

We next tested whether pharmacological blockade of KCa3.1 functions can suppress this phenotype and treated the mice with Senicapoc-containing chow at a dose of 30mg/kg/day [44] during the 2-weeks of DOX treatment.

The Senicapoc treatment gave rise to total plasma concentrations of 254+/−61 nM (n=11) and tissue levels in the skin of 2.9+/−0.9 μM (n=4) at the time of sacrifice. At these total concentrations (and assuming a protein binding of 90%), pharmacological inhibition of the channel can be expected because concentrations of Senicapoc in the skin are at least 2 orders of magnitude above the reported IC_50_ of 10 nM [12, 37].

As shown in Figure 5 A, we report that the Senicapoc treatment did not prevent the pathological skin alterations, but significantly reduced intra-epidermal edema by ≈60%, hyperplasia by ≈50%, as well as a trend in reduction of hyperkeratosis and fibrosis.

**Figure 5:**
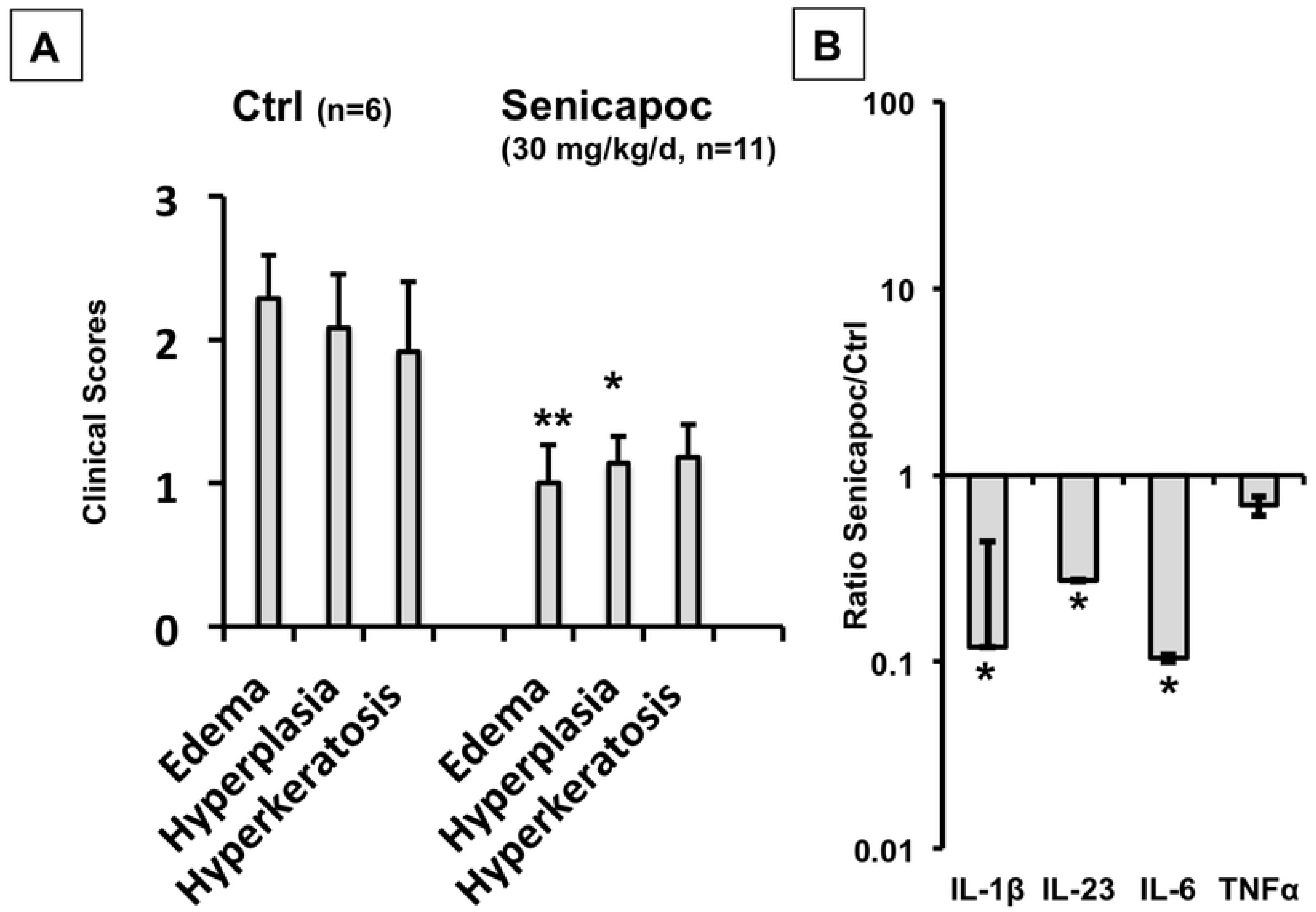
A) Senicapoc suppressed skin pathology and induction of IL-β1, IL-6, IL-23 in DOX-treated KCa3.1+ mice. B) Data (Senicapoc/Ctrl) are given as means +/− SEM, n=5 each, *P<0.05, **P<0.01 Student’s T test.

Overall, this Senicapoc trial suggests that the skin pathology was caused largely by the induced function of KCa3.1 and demonstrates that Senicapoc is able to reduce the earliest strong alteration, i.e. intra-epidermal edema (Fig 3 C and D), presumably by blocking KCa.3.1-induced disturbances of epidermal water-salt homeostasis.

Considering cytokine levels in the skin (Fig 5 B), Senicapoc significantly reduced mRNA-expression levels of IL-β1 (≈−88%), IL-23 (≈−77%), as well as of IL-6 (≈−90%) when compared to DOX-treated control animals, while TNFα mRNA-expression levels were unchanged. These lower levels of pro-proliferative cytokines could explain the lower degree of hyperplasia.

## DISCUSSION

The present study introduces a genetic model of epidermal KCa3.1-induction to investigate the physiological and potential pathophysiological significance of this channel capable of controlling cell volume, growth, and cellular salt/water homeostasis in the epidermis. In fact, it is the first study describing the consequences of genetically encoded induction of an ion channel above basal levels in the skin. Our results demonstrate that KCa3.1 over-expression was capable of causing a severe disturbance of epidermal homeostasis.

This is based on the following lines of evidence: 1) KCa3.1-induction produced intra-epidermal edema (spongiosis). 2) KCa3.1-induction produced progressive epidermal hyperplasia and hyperkeratosis causing severe itch and ulcers. Finally, 3) KCa3.1-induction strongly up-regulated epidermal (i.e., keratinocyte) synthesis of the pro-proliferative, auto-stimulatory, and pro-inflammatory cytokines, in particular, IL-β1, IL-23, and TNFα. Taken together, KCa3.1-induction in the epidermis produced skin pathology in mice that resembled the pathological features of itchy eczematous dermatitis [42].

The physiological and pathophysiological significance of KCa3.1 in the skin has been elusive until now. So far, keratinocyte KCa3.1 has been studied – superficially though - by mRNA-expression experiments in cultured human keratinocytes [5] and immune histochemistry in rat epidermis [39], which did not provide much information about the role of the channel in epidermal homeostasis *in-vivo*. Here we intended to provide new knowledge by generating a murine model of genetic conditional induction of a murine KCa3.1-transgene in epidermis (Fig 1).

The KCa3.1 induction in DOX-treated animals was specific for skin epidermis as concluded from 800-fold overexpression. Concerning other tissues, we found a much less pronounced 20-fold overexpression in the intestine (see below).

We confirmed KCa3.1 transgene function after *in-vivo* DOX-treatment by patch-clamp electrophysiology on isolated keratinocyte, which unequivocally revealed large KCa3.1-currents being sensitive to a KCa3.1-inhibitor displaying the biophysical and pharmacological characteristics of KCa3.1 (Fig 1 C) [12]. In keratinocytes from untreated mice, KCa3.1 currents were very small or undetectable suggesting low basal channel expression or a few KCa3.1-expressing cells. IHC demonstrated appreciable immune reactivity in the hyperplastic epidermis of the DOX-treated mice. Immune reactivity was also found in the thin epidermis of untreated mice, suggesting some constitutive protein expression in addition to mRNA-expression of KCa3.1 (Fig 1 D).

A major outcome of our study was that KCa3.1-induction produced visibly piloerection, scaly skin patches, and intense scratching behavior that in turn gave rise to bloody ulcerative lesions (Fig 2 and S1 Movie). These symptoms are similar to those of itchy eczematous dermatitis in humans [42]. In analysis of histological sections, we observed intra-epidermal edema (sub-acute spongiosis) and ensuing progressive hyperplasia, and hyperkeratosis as histological features of chronic eczema [42].

It is also worth mentioning that sub-chronic conditional gene deletion of KCa3.1 in the epidermis as well as life-long KCa3.1-deficiency [16] complete did not produce any skin alterations, indicating that basal KCa3.1 expression is apparently not crucial for epidermal homeostasis. In keeping with the induction of KCa3.1 in the small intestine, we also demonstrate a mild intestinal phenotype characterized by moderate chyme accumulation and lower propulsive spontaneous motility, which is the content of a separate report [45].

Concerning the cellular mechanisms, by which KCa3.1-induction produced this skin condition, the morphological and molecular biological alterations described here agree well with the known physiological functions of KCa3.1 [12]: 1) The intra-epidermal edema can be explained by KCa3.1’s ability to move K^+^ and concomitantly water and Cl^−^ into the extracellular compartment [13, 15, 20]. Overexpression of the channel could do this in an excessive manner leading to the observed expansion of the extracellular compartment and/or cell shrinkage producing intracellular gaps. In fact, intra-epidermal edema was already pronounced at the early time point (1 week), when hyperplasia and hyperkeratosis was still in an initial phase. Intra-epidermal edema was of similar grade later, when hyperplasia and hyperkeratosis progressed further.

Therefore, intra-epidermal edema can be considered the starting point for the latter alterations.

2) In addition to its function as a K^+^ secreting channel, KCa3.1 activity is known to produce strong membrane hyperpolarization, which in turn potentiates calcium-influx enabling long-lasting elevations in [Ca^2+^]_I_ (for review see: [1, 36]). In several cell systems, this elevation of [Ca^2+^]_I_ has been shown to be required for the sustained initiation of several cellular processes, including proliferation and migration [1, 46].

KCa3.1-mediated hyperpolarization and ensuing amplification of [Ca^2+^]_I_ signaling is known to regulate cytokine production in T cells, activated fibroblasts or smooth muscle cells (for review see: [1]). Accordingly, inhibition or genetic knockdown of the channel has been reported to reduce IL-2 production by T cells [46–47] and IL-β1 and TNFα levels by microglia in ischemic stroke [19]. Here, we showed that the reverse maneuver (induction) resulted in a higher expression of IL-β1, IL-23, and TNFα in keratinocytes of the epidermis (Fig 5). This keratinocyte cytokine production may drive hyperplasia and hyperkeratosis in an auto-stimulatory fashion.

Considering the impressive magnitude of hyperplasia and hyperkeratosis (Figure 3), we speculate that this is a secondary and overshooting repair mechanism in response to intra-epidermal edema and epidermal destabilization.

In summary, we provide first mechanistic evidence that KCa3.1 induction produced skin pathology *in-vivo* by causing extracellular fluid accumulation and epidermal destabilization as a primary event and secondary phenotypic switch to a proliferative keratinocyte phenotype (PCNA-high, Fig 3E), producing epidermal hyperplasia.

It is worth mentioning that the selective KCa3.1-blocker Senicapoc significantly reduced both the primary intra-epidermal edema and the subsequent hyperplasia (Fig 5). The reduction in edema is somewhat reminiscent of the reported reduction in *in-vitro* cyst formation by kidney cells from patients with autosomal-dominant polycystic kidney disease with a KCa3.1 blocker [25]. Moreover, Senicapoc treatment reduced IL-β1, IL-6, and IL-23, and TNFα mRNA-expression levels in the affected skin (Fig 5). These data strongly indicate that these morphological alterations and the higher expression pro-proliferative cytokines as mechanistic drivers of epidermal hyperplasia are mediated by channel functions. Because the treatment did not fully suppress the phenotype, we cannot fully exclude additional local or systemic mechanisms.

Compared with human skin diseases, the phenotype in KCa3.1+ mice is strikingly similar to eczematous dermatitis with spongiosis, epidermal hyperplasia, and itch, as important pathological features [42]. Yet, the cellular mechanisms causing this condition and, particularly, spongiosis are poorly understood. In this regard, our findings suggest that epidermal KCa3.1 could be a mechanistic player. At present a pathomechanistic role of KCa3.1 has not been shown for human skin conditions with the exception of role in melanoma cell proliferation. Yet, our study provides the rationale to investigate KCa3.1-function in specifically eczematous dermatitis with keratinocyte hyperplasia and in other skin pathologies, characterized by excessive keratinocyte growth such as psoriasis, and test clinical efficacy of KCa3.1-inhibitors.

In conclusion, epidermal KCa3.1 overexpression in murine skin produces itchy eczematous dermatitis. This can be dampened by pharmacological channel inhibition. Future target identification and validation studies in patients will show whether KCa3.1-inhibitors are of therapeutic utility in human skin pathologies.

## CONFLICT OF INTEREST

The authors state no conflict of interest.

## ACKNOWLEDGEMENTS

We thank Andrea Lorda for excellent technical assistance and the scientific staff of the IACS: Dr. Maria Royo (IACS), Dr. Mark Strunk, Alicia de Diego Olmos and the staff of animal of the animal core facility IACS for IHC stains, genotyping, animal breeding and collecting tissue, respectively.

## Supporting Information

1) S1 Fig. Supplemental Data (multi-paneled figure)

2) S1 Movie. Supplemental Media file 1; Description: Two sequences showing mice treated with DOX for two weeks and a sequence showing control mice receiving only sucrose. Of the last two sequences the first shows a mouse from the Senicapoc trial that did not receive Senicapoc. The second sequence shows a mouse that received Senicapoc.

3) Data files:

3.1 Data qRTPCR

3.2 Scores (skin pathology) and electrophysiological data

## Abbreviations

CASP3: Caspase-3
DOX: Doxycycline
KCa3.1: intermediate-conductance calcium-activated potassium channel
PCNA: proliferating cell nuclear antigen

## Notes

Funding sources: This work was supported by the Government of Aragón (DGA-METIC–consortium; B04_17R), FP7-PEOPLE-MC-CIG to RK, FIS-CB06/07/1036 to PG and RK, ERC-2014-StG-638284, DPI2016-75458-R, T39_17R to EP; FIS-PI16/02112 to ALGO; NHLBI-R01-HL080173, NCRR-P20-RR018751 to HM. JLG received a DGA-PhD scholarship (C072/2014). BMB was supported by the NCATS (UL1 TR001860 and linked award TL1 TR001861). KLH received sabbatical support from the University of Otago.

